# Previous oil exposure alters oil avoidance behavior in a common marsh fish, the Gulf Killifish *Fundulus grandis*

**DOI:** 10.1101/2020.06.15.152413

**Authors:** Charles W. Martin, Ashley M. McDonald, Guillaume Rieucau, Brian J. Roberts

## Abstract

Oil spills threaten the structure and function of ecological communities. In the northern Gulf of Mexico, the 2010 Deepwater Horizon spill was among the largest marine disasters in history. While many predicted catastrophic consequences for nearshore fishes, field studies indicate surprising resilience in populations and communities. One potential mechanism for this resilience is the recognition and behavioral avoidance at small spatial scales of the toxic chemical constituents found in oil. Previous research indicates many marsh fishes have the capacity to avoid oil contaminated areas. Here, we test whether prior oil exposure of a common marsh fish, the Gulf killifish *Fundulus grandis*, alters this avoidance response. Using choice tests between unoiled and a range of oiled sediments, we found that, even at low levels of previous exposure, killifish lose recognition of oiled sediments. Preference for unoiled sediments was lost across the entire range of oil concentrations tested here after oil exposure, and some evidence for preference of oiled sediments was even demonstrated. These results provide evidence for lack of response to toxic environments in exposed individuals, suggesting sublethal impairment of sensory mechanisms on an individual level despite organism survival. Future research should highlight additional sublethal consequences that affect ecosystem and food web functioning.

## Introduction

The 2010 Deepwater Horizon (DwH) oil spill in the Gulf of Mexico (USA) impacted nearshore ecosystems from the Louisiana coast to the Florida Panhandle (Michel et al. 2013). Over 87 days, approximately 3.19 million barrels of oil (US vs. BP 2015) ultimately covered an estimated 180,000 km^2^ of Gulf waters and ∼1100 km of coastal wetland shorelines with ∼95% of these being in Louisiana (Mitra et al. 2012, Michel et al. 2013, Nixon et al. 2016). The spill predominantly impacted salt marshes in nearshore areas, with significant economic implications given they serve as key habitats for the young of many commercial and recreational fishery species (Peterson and Turner 1994, Lellis-Dibble et al. 2008, Rozas et al. 2013). To date, field assessments of coastal fish populations and communities have shown resistance and resilience to, and in some cases rapid recovery from, the toxic effects of oil (Moody et al. 2013, Fodrie et al. 2014, Able et al. 2015, Schaefer et al. 2016). Several potential explanations have been given for the unanticipated lack of severe impacts to populations/communities in nearshore ecosystems, including behavioral emigration from oiled areas, sublethal impacts to individuals (Whitehead et al. 2012, Dubansky et al. 2013, Martin 2017a) that do not translate to higher levels of organization, indirect food web mechanisms that provide predator release and/or stimulation of production, and cessation of fishing (Fodrie et al. 2014, Schaefer et al. 2016, Martin et al. 2020).

Studies across a wide range of organisms, from zooplankton (Seuront 2010) to marine mammals (Smultea and Würsing 1995, Ackleh et al. 2012), have indicated complex recognition patterns and behavioral avoidance of oiled conditions. For example, calanoid copepods alter swimming behavior to avoid water-soluble diesel oil to limit exposure (Seuront 2010). At larger spatial scales, sperm whales relocated from their historically occupied areas due to the DwH oil spill (Ackleh et al. 2012). Dolphins exhibit similar avoidance responses (Smultea and Würsing 1995) and have been trained in the detection of oil (Geraci et al. 1983). Conversely, American lobsters have shown attraction to hydrocarbons such as kerosene (Atema 1976), which has been used as a bait by commercial fishermen.

Fishes have strong behavioral responses to crude oil contamination (Weis et al. 2001, Martin 2017a, Schlenker 2019a). Freshwater fishes such as fathead minnows *Pimephales promelas* (Farr et al. 1995), rainbow trout *Oncorhynchus mykiss* (Carr et al. 1990), pink salmon fry *Oncorhynchus gorbuscha* (Rice 1973), Caspian roach *Rutilus caspicus* (Lari et al. 2015), and striped bass *Morone saxatilis* (Carr et al. 1990) avoid hydrocarbon contaminated areas, albeit at different thresholds and concentrations. Estuarine and marine fishes such as flatfishes (Moles et al. 1994), juvenile spot *Leiostomus xanthurus* (Hinkle-Conn et al. 1998), European seabass *Dicentrarchus labrax* (Claireaux et al. 2018), and mahi-mahi *Coryphaena hippurus* (Schlenker 2019a) all respond behaviorally to avoid the toxic chemical contaminants found in petroleum hydrocarbons.

In salt marshes, previous work (Martin 2017a) has demonstrated that key fishes such as Gulf killifish *Fundulus grandis*, sailfin molly *Poecilia latipinna*, and sheepshead minnow *Cyprinodon variegatus* avoid oil contaminated sediments, but display a reduced response to weathered oil indicating they likely react to the volatile, aromatic compounds that are lost as oil degrades due to a combination of factors including UV exposure, wave action, microbial processing, among others. These shallow, soft-sediment areas dominate the inshore reaches of the northern Gulf of Mexico (Connor and Day 1987) and benthic organisms living within and on these sediments serve as important food sources for numerous species, including one of the most abundant Gulf coast marsh species *F. grandis*. For example, Rozas and LaSalle (1990) reported that the major dietary constituents of *F. grandis* are found associated with these sediments: fiddler crabs, amphipods, and hydrobiid snails. As such, recognition of the quality and contamination of these habitats is critical for *F. grandis* as these areas are linked with successful foraging and fitness.

Previous research indicates that oil exposure can inhibit or damage sensory mechanisms used by fishes to detect oil. For example, exposure to the water-accommodated fraction of oil in Atlantic stingrays *Hypanus sabinus* damaged olfactory function (Cave et al. 2018). After oil exposure, bicolor damselfish *Stegastes partitus* showed a reduction in the response to conspecific alarm cues (Schlenker et al. 2019b). This damage to olfactory mechanisms or central nervous processing may further reduce organism’s capacity to detect and respond to oil contamination (sensu Schlenker et al. 2019a). Here, we present the result of a series of experiments to test the effects of previous exposure on oil-contaminated sediment avoidance in salt marsh fish. Using the Gulf killifish *F. grandis*, a widely-used sentinel species for toxicological and ecological studies (Able et al. 2015, Vastano et al. 2017, Jensen et al. 2019), we exposed animals to a range of experimentally oiled marsh conditions and then subsequently tested behavioral avoidance patterns in simple choice tests. The overarching objective of this project is to determine if prior exposure influences avoidance behavior and, if so, to identify what exposure level influences these changes.

## Methods

### Fish Exposure

Fish were exposed to oil for 10-15 days in experimentally oiled marsh mesocosms at the Louisiana Universities Marine Consortium in Cocodrie, LA during August/September 2019. Briefly, we utilized 12 hydrologically independent *Spartina alterniflora* marsh mesocosms (3.05m diameter, 1.83m height) each with its own paired tidal surge tank generating daily tidal cycles with range of 25cm (flooding marsh ∼10cm at high tide) via a water control system of blowers and airlifts. Light Louisiana Sweet (LLS) blended crude oil, API Gravity 40.1, similar to that which was released in the Deepwater Horizon oil spill, was acquired from Placid Refining Company LLC in May 2018. This oil was evaporatively weathered by 30% of its volatile components, as measured by gas chromatography, using a nitrogen gas sparging system placed in the barrel of liquid oil as received from Placid over a period of 150 days. The bubbling system not only expedited evaporation, but also mixed the contents of the barrel to ensure conformity of its contents. On July 8, 2019, weathered oil was applied to each tank at high tide at one of four concentrations, scaling roughly to SCAT categories observed on shorelines after the Deepwater Horizon spill (Michel et al. 2013, Nixon et al. 2016): control/no oil (0.0 L oil m^-2^), low (0.1 L oil m^-2^), medium (0.5 L oil m^-2^), and high (3.0 L oil m^-2^) oil concentrations.

We captured *F. grandis* from the nearby salt marsh using baited minnow traps and held in separate aquaria for 2-4 days to minimize any mortality due to handling before being introduced to the 15 cm wide trough surrounding plants within each mesocosm. During low tide, fish were restricted to a 15cm deep water column and at high tide events, fish gained access to oiled or unoiled experimental marsh platforms (at a water depth of ∼10cm) to forage. A total of 18 fish were added to each tank between 22 August (12 fish added) and 27 August (6 additional fish) 2019. A total of 54 fish/treatment were introduced to mesocosms and recaptured prior to experiments using dip nets (with the treatment exposed period being between 45 and 60 days after initial oiling of the mesocosms). The surface (0-5cm) total petroleum hydrocarbon (TPH) concentrations in the high oil treatments (419 + 4 mg / g soil) were ∼10 and ∼40 times higher than in the moderate (39 ± 6 mg / g soil) and low (10 ± 0.4 mg / g soil) oil treatments (mean of 19 August and 9 September samplings; Overton et al. 2020). We noted some mortality and lack of recapture for some treatments, particularly in medium and high oil mesocosms, which precluded the full range of preference tests in the avoidance experiment (described below). Specifically, out of the original 54 fish released we recaptured 54, 34, 24, and 10 fish in control, low, medium, and high treatments, respectively. These differences in mortality corresponded to survivorship of 100% (control), 62.0% (low), 44.4% (medium), and 18.5% (high) in mesocosm treatments.

### Avoidance Experiment

To test the behavioral response of fishes with varying oil exposure histories to different oil concentrations, we used a choice test following the design reported in Martin (2017a). Thirty-eight-liter aquaria were filled with 3 L total sediment to a depth of 18 cm. Each aquarium offered a choice between unoiled and oiled sediment, randomized on each side of the tank. Concentrations followed previous experiments (Martin et al. 2015, Martin 2017a), and fish were given a choice between no oil and low (10 mL oil L^-1^ sediment), medium (20 mL oil L^-1^ sediment), or high (40 mL oil L^-1^ sediment) contamination. In these tests, we used 25% weathered oil (from the same source barrel as used in the larger mesocosm experiment) as this was representative of what came inshore (Reddy et al. 2012) and is the same degree of weathering used in previous experiments (Martin 2017a). We mixed oil with sediment on the randomized side of the tank at the assigned concentration and a thin layer (approximately 2 cm deep) of unoiled sand placed on top to prevent the oil/sand mixture from floating after filling tanks with water and affecting the adjacent, unoiled side of the tank. From an experimental design perspective, using unstructured sediment as choices prevents any confounding preference due to structural refuge using other habitats such as marsh grass or submerged vegetation.

All *F. grandis* used in this study were adult individuals between 57 mm and 105 mm and used only once in a trial. Mortality during exposure limited the number of available fish, and as a result we replicated most comparisons 8 times, except for medium exposure fish (no oil vs medium oil was replicated 4 times) and high exposure fish (only no vs low oil was tested with 8 replicates). We held salinity constant at 7.0 and temperature ranged from 26.8-28.6°C during trials (both comparable to the conditions in the mesocosms at the time of collection).

For each trial, one fish from a randomized exposure was introduced to the tank, allowed a 5-minute acclimation period, and its movements between the two sides of the tank recorded using a GoPro camera over the course of 10 minutes. This trial period mimics previous fish behavioral experiments (Gerlach et al. 2007, Paris et al. 2013, Martin 2017a). The side of the tank occupied by fish (no oil or assigned oil treatment) was recorded by analyzing a frame every 30 seconds and noting fish position within the tank. The proportion of time in each side was then calculated as the number of observations taken on that side divided by the total observations. To graphically display data, a ratio of the number of times fish occupied each side of the tank was generated and plotted, such that deviation below 1 indicates avoidance of oil and above the line denotes preference for oil.

Data were statistically analyzed in 2 ways: 1) to determine whether fish deviated from an expected 1:1 occupancy pattern, we conducted a paired t-test for each previous exposure level and oil vs no oil comparison (Peterson and Renaud 1989) and 2) to compare differences across treatments, we analyzed proportion of time spent in oil using a general linear model (GLM) with factors of previous exposure and oil vs. no oil comparison. Tukey’s post hoc test was used to determine significant pairwise differences. Assumptions (normality and homogeneity of variance) were tested for all comparisons and nonparametric alternatives (signed rank test) used if transformations failed to meet assumptions and considered results significant at p<0.05.

## Results

Previous exposure influenced fish preference patterns for oil contaminated sediments (Figure 1, Table 1). Unexposed (control) fish significantly avoided the oiled side of aquaria, regardless of the oil concentration choice given (Table 1). After exposure to oil, even at low concentrations, this avoidance response disappeared. Fish unexposed to oil spent on average 66% of time over uncontaminated sediments, a trend that decreased with previous exposure to low (52%), medium (55%), and high (44%) concentrations. At high previous exposure concentrations, we noted that many fish spent more time over oiled sediments in the aquaria, although not significant (Table 1). We visually inspected the data and noted one trial where behavior differed from others. In this case the fish spent 65% over unoiled sediments. After removing this trial and reanalyzing data, high exposure led to significant (t(6)=2.517; p=0.045) preference for oiled sediments at low concentration over unoiled sediments. Note that Figure 1 only reports this analysis with the fully dataset (some treatments were unable to be conducted due to lack of experimental organisms, as noted above).

**Table 1.** Statistical results for each previous exposure and comparison. Control fish preferred no oil and the significant high exposure fish (with outlier removed) preferred oiled sediments. Significant p-values are shown in bold.

| Previous Exposure | Comparison | Statistical Test | N | Test Statistic* | p |
| --- | --- | --- | --- | --- | --- |
| Control (No Oil) | Low Oil vs No Oil | Paired t-test | 8 | -4.204 | **0.004** |
|  | Medium Oil vs No Oil | Paired t-test | 8 | -7.105 | **<0.001** |
|  | High Oil vs No Oil | Signed rank | 8 | 2.388 | **0.016** |
| Low Oil | Low Oil vs No Oil | Paired t-test | 8 | <0.001 | 1.000 |
|  | Medium Oil vs No Oil | Paired t-test | 8 | 0.235 | 0.821 |
|  | High Oil vs No Oil | Signed rank | 8 | 1.550 | 0.148 |
| Medium Oil | Low Oil vs No Oil | Paired t-test | 8 | -1.546 | 0.166 |
|  | Medium Oil vs No Oil | Paired t-test | 4 | 0.378 | 0.731 |
|  | High Oil vs No Oil | Paired t-test | 8 | -1.080 | 0.316 |
| High Oil | Low Oil vs No Oil | Paired t-test | 8 | 1.418 | 0.199 |
| High Oil (no outlier) | Low Oil vs No Oil | Paired t-test | 7 | 2.517 | **0.045** |

\*Test statistic: T for Paired t-test, Z for Signed Rank test

**Figure 1.**
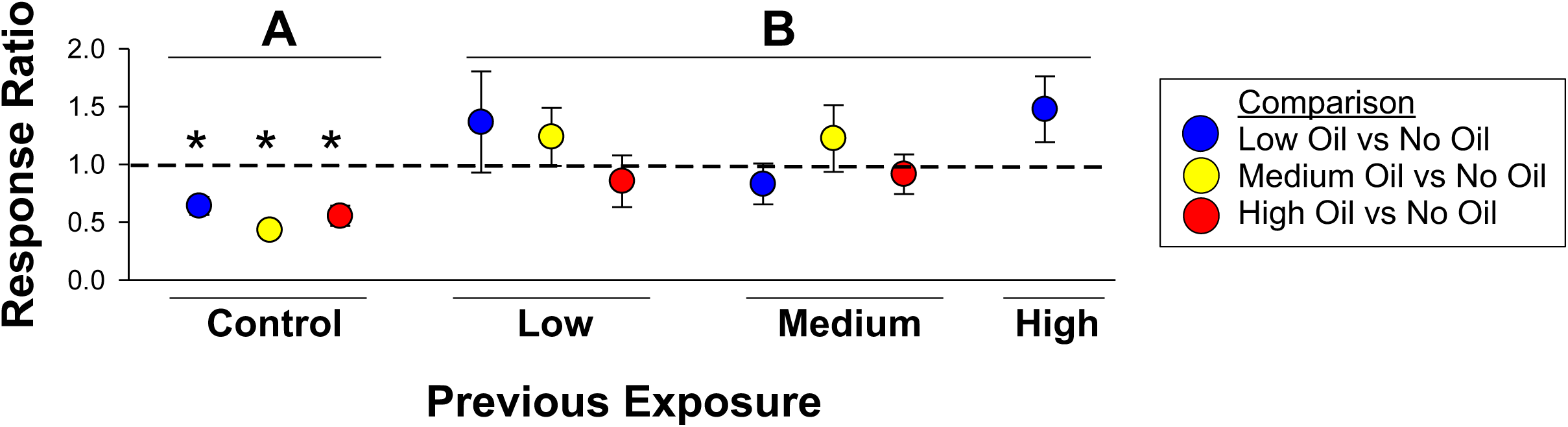
NOTE may have to chance to white/gray/black color scheme depending on journal. Ratio of time spent in oil to time spent in no oil side of tank across previous exposures with the dashed line indicating 1:1 (no preference). Colors represent values for preference comparisons between no oil and low (blue), medium (yellow), and high (red) oil concentrations. Asterisks indicate fish deviate from an expected 1:1 occupancy pattern (Table 1) and different letters indicate statistical differences among treatments. Note this figure reports analyses only on the full dataset.

The oil concentration used in the choice test had a smaller influence on fish behavior. Results of the GLM confirmed these results, which indicated that previous exposure (F_3,70_=6.83, p<0.001), not oil concentration in preference test (F_2,70_=0.38, p=0.684), drove behavioral patterns. Pairwise comparisons indicated that control, unexposed fish had significantly stronger avoidance of oil than exposures at low (p=0.0034), medium (p=0.0337), or high (p=0.0029) exposure, with low, medium, and high not significantly different from each other (p>0.05).

## Discussion

The 2010 DwH oil spill was an unprecedented stressor for northern Gulf ecosystems, with oil impacting emergent and submerged plants (Lin and Mendelssohn 2012, Silliman et al. 2012), invertebrates (McCall and Pennings 2012, Powers et al. 2017, Zengel et al. 2016), and fishes (Fodrie and Heck 2011, Able et al. 2015, Schaefer et al. 2016). While the full effect on the nearshore food web may not yet be fully realized because of the numerous indirect, food web mechanisms (McCann et al. 2017), many studies have documented significant impacts to molecular, genomics, and development of fishes (Whitehead et al. 2012, Dubansky et al. 2013) and resilience in populations and communities to oil’s toxic effects (Fodrie et al. 2014, Martin et al. 2020). Among the proposed explanations for this resilience, despite oil’s known toxicity, is the behavioral emigration of organisms at small spatial scales to avoid exposure to contamination (Martin 2017a). Here, we demonstrate that exposure to oiled marshes, even at low concentrations of 0.1 L oil m^-2^, can impact a common marsh fish’s ability to detect and avoid oil contaminated sediments.

After removal of one outlier, exposure to oil at high concentrations resulted in preference for oiled sediments in the choice test. We hypothesize that this could be due to relaxation of normal physiological functioning causing an anaesthetic effect (Barron et al. 2004). Several studies have documented a narcotic effect on fishes due to chronic exposure to oil (Barron et al. 2004, Incardona et al. 2004). This can result in a nervous system sedation (Lin and Tjeerdema 2008), as well as increased respiratory rate (Brocksen and Bailey 1973) and decreased swimming performance (Stieglitz et al. 2016). These effects can be reversed, however, usually on the order of days after exposure has been removed (Brocksen and Bailey 1973). In this experiment, we only conducted choice trials at low concentrations for the high exposure treatments, leaving open the possibility that a stronger response could have been observed had more fish survived the exposure.

Previous exposure to oil is known to have many negative consequences for individuals. For example, unexposed mahi-mahi avoided higher concentrations of water accommodated fraction of oil, but exposed individuals demonstrated a lack of response (Schlenker et al. 2019a). Moreover, damage to the higher order central nervous system processing was implicated in this decreased oil avoidance behavior. Olfactory damage is also known to occur with oil exposure in some fishes (such as Atlantic stingrays, Cave et al. 2018) and reduced recognition of threats via these cues is also possible (Schlenker et al. 2019b). Whether highly weathered crude oil elicits similar effects and the specific chemical constituents in oil involved in these detrimental physiological changes remain unclear. Given that weathered oil comprised the bulk of the oil that came ashore (Reddy et al. 2012), it is likely that fish survival may have been higher than expected and thus sublethal effects may constitute the largest impact on marsh organisms. In our exposures, we observed mortality at all oil levels, especially at high oil concentrations, with some survival and these findings support the notion that even if fish survive there are significant behavioral responses that might influence their long term survival.

Given the importance of olfaction for many critical activities, such as foraging (Webster et al. 2007, Johannesen et al. 2012), habitat recognition (Forward et al. 2003, Benfield and Aldrich 1992, Martin 2017b), and predator avoidance (Dixson et al. 2010, Martin et al. 2010, Martin 2014), it is possible that these and other sublethal effects may have great consequences that, to date, have remained largely unexplored in the wake of the DwH oil spill. Previous studies have indicated that oil can have other important sublethal effects on fishes and invertebrates. For example, oil presence triggered a 60% decrease in penaeid shrimp *Farfantepenaeus aztecus* growth rate (Rozas et al. 2014) and foraging by darter gobies *Gobionellus boleosoma* can change 50-100% in sediments highly contaminated with diesel fuel (Gregg et al. 1997). Spot *L. xanthurus* reduce feeding strikes in the presence of diesel oil (Hinkle-Conn et al. 1998). Given the known deleterious impacts to other fishes, we anticipate that similar sublethal consequences were present in marsh fishes after DwH, but have remained understudied. We propose that additional research on the sublethal effects of oil need to be conducted to gain a broader understanding of the full scope of DwH damages to northern Gulf of Mexico ecosystems.

Oil released from the DwH drilling rig was burned at the surface, collected on the water or as it came ashore on wetlands and beaches, and chemically dispersed using Corexit® dispersant (Peterson et al. 2012, Michel et al. 2013, Nixon et al. 2016). Wetlands accounted for over half of the oiled shoreline (∼1100 of ∼2100km), with > 95% of oiled marshes in Louisiana (Nixon et al. 2016). However, much of the oil remains unaccounted for (McNutt et al. 2012) and is thought to reside in sediments throughout the region. Previous spills such as the *Exxon Valdez* (Renner et al 2006, Li and Boufadel 2010), *Florida* barge in Massachusetts (Culbertson et al. 2008), and *Ixtoc-I* (Schrope 2010) all indicate that oil can persist buried in the sediment where oil weathering rates are low (Boufadel et al. 2010). Thus, marsh fishes may be vulnerable to sublethal oil exposure and the loss of avoidance behaviors in marsh species may have more subtle, but still substantial, implications for the marsh food web long after the oiling event.

Unlike in pelagic species where exposure is comparatively more limited because oil moves long distances across the surface with currents and wind, weathers, biodegrades, or sinks to deep sediments, once oil reaches marsh sediments exposure may be extended. Once exposed, these marsh fishes lose recognition and remain vulnerable to oil contamination in the short or long term as oil gets trapped by the plants and buried or slowly degraded over time. Enhancement of erosion rates (Silliman et al. 2012, Martin et al. 2015, Turner et al. 2016) and sediment remobilization after large storm events, such as the frequent Gulf of Mexico hurricanes (Khanna et al. 2013, Michel et al. 2013), may re-expose remaining oil to saltmarsh flora and fauna, continuing to sublethally impact organisms for decades to come. Many resident and transient species spend some part of their life cycles in these contaminated areas and could be impacted for sustained periods, necessitating the need for continued study of oil impacts in these vital ecosystems.

## Acknowledgements

This research was made possible by a grant from The Gulf of Mexico Research Initiative. The funders had no role in the design, execution, or analyses of this project. Data are publicly available through the Gulf of Mexico Research Initiative Information & Data Cooperative (GRIIDC) at https://data.gulfresearchinitiative.org (doi:<10.7266/n7-3qjh-2f11>). We thank the staff at Louisiana Universities Marine Consortium for the facilities and logistical support needed to make this project possible, particularly members of the Roberts lab for the initiation and maintenance of the mesocosm experiment (Charles Schutte, Ryann Rossi, Ekaterina Bulygina, Stephanie Plaisance, Caitlin Bauer) and members of the Education and Outreach team (Murt Conover, Aaron Bacala, and Tori Lambert) that helped in organism collection and holding.

## Notes

### Competing Interest Statement

The authors have declared no competing interest.

